# Self-organizing motors divide active liquid droplets

**DOI:** 10.1101/403121

**Authors:** Kimberly L. Weirich, Kinjal Dasbiswas, Thomas A. Witten, Suriyanarayanan Vaikuntanathan, Margaret L. Gardel

**Affiliations:** James Franck Institute, University of Chicago, Chicago, IL 60637; Department of Physics, University of Chicago, Chicago, IL 60637; Department of Chemistry, University of Chicago, Chicago, IL 60637

## Abstract

The cytoskeleton is a collection of protein assemblies that dynamically impose spatial structure in cells and coordinate processes such as cell division and mechanical regulation. Biopolymer filaments, cross-linking proteins, and enzymatically active motor proteins collectively self-organize into various precise cytoskeletal assemblies critical for specific biological functions. An outstanding question is how the precise spatial organization arises from the component macromolecules. We develop a new system to investigate simple physical mechanisms of self-organization in biological assemblies. Using a minimal set of purified proteins, we create droplets of cross-linked biopolymer filaments. Through the addition of enzymatically active motor proteins we construct composite assemblies, evocative of cellular structures such as spindles, where the inherent anisotropy drives motor self-organization and droplet deformation. These results suggest that simple physical principles underlie the self-organization in complex biological assemblies and inform bio-inspired materials design.

## Introduction

Spontaneous organization occurs in biology at all length scales, from organelles and tissues to organisms and populations (1-3). For instance, in cell division self-organized macromolecular assemblies drive localization of chromosomes to the cell midplane, segregate chromosomes, and split the cell into two daughter cells (4). Each stage relies on molecular species within the cytoplasm precisely spatially and temporally localizing to regulate biochemical signals and generate mechanical stresses. While molecular self-organization principles have been intensely studied (3), we still lack understanding needed to reconstruct these intricate processes from purified components *in vitro*. Elucidating such principles would yield insights into *in vivo* regulation as well as inform the design of novel, bio-inspired, soft materials (5-8).

Biological molecules form unusual and adaptive machinery that drive physiological processes. The cytoskeleton, composed of biopolymer filaments and associated binding partners, constitutes a broad class of assemblies that regulate diverse processes ranging from cell motility and division to intracellular transport (9). A critical component of the cytoskeleton is actin, a protein which polymerize into semi-flexible filaments that assemble into elastic-like meshworks and bundles (10). These actin assemblies are responsible for regulating cell shape changes and mechanics (10). Myosin II motors, found as bipolar filaments of 10-100 motors, drive relative motion of actin filaments, resulting in force generation (11). When reconstituted *in vitro*, assemblies of actomyosin exhibit striking dynamical behavior, where myosin motors drive remodeling of actin networks, resulting in network compaction and remodeling (12).

Besides the cytoskeleton, self-organized assemblies in the cytoplasm are ubiquitous. Liquid-like phases have been recently identified in a variety of proteins and nucleic acids (13, 14) and are thought to play important roles in subcellular compartmentalization (15). Macromolecular liquid droplets have also been proposed to have been the earliest primitive cells, or protocells, where reactive contents phase separated from the surrounding environment (16). These macromolecular droplets are thought of as isotropic liquids (13), characterized by a single viscosity and surface tension and lacking internal structure. We recently reported another type of macromolecular droplet, where the actin cross-linker, filamin, condenses short actin filaments into liquid droplets (17). While actin filament-based liquids have characteristics of conventional liquids, they are anisotropic due to internal structure from densely packed filaments, so instead form liquid crystals (17-19). The internal liquid structure of molecular liquid crystals can be harnessed to drive processes such as the spatial localization of colloidal particles (20, 21). Although the effect of active stresses generated by molecular motors in nematics is an emerging area of research (22-25), in droplets it is unknown and has no analog in experimental molecular liquid crystal systems.

Here, we report macromolecular droplets, composed of actin liquid crystal embedded with the molecular motor myosin II, that robustly self-organize and spontaneously change shape. Intriguingly, these droplets exhibit two essential features of cell division: localization of species at the droplet midplane and droplet bisection into two “daughter” droplets. Using fluorescence microscopy, we find that individual myosin II filaments dispersed throughout the actin droplet migrate to the droplet midplane. Active myosin filaments form clusters that locally distort actin alignment and, when confined to droplets, drive droplet division into two daughter droplets of equal volume. We develop a continuum model, based on the alignment of rods in a structured fluid to describe motor centering. By extending the model, through representing a motor cluster as a spherical colloid which favors radial alignment of actin, we capture droplet division. Of note, the role of enzymatic activity is to change the geometry of local actomyosin interactions and is captured by a free energy model by considering changes in motor geometry. Our results highlight structural changes to droplets induced by colloidal objects, which potentially provides insight into physical mechanisms of center finding in biology and cell division, as well as informs the design of novel geometry-sensing and adaptive materials.

## Results

To create structured actin droplets, we cross-link dilute (2.6 μM actin monomer), short (∼ 200 nm) actin filaments with the actin-binding protein, filamin (10 mol%, SI). Using confocal microscopy, we find that filamin quickly condenses short actin filaments into micron-sized spindle-shaped droplets, called tactoids (17). These droplets are composed of densely packed actin filaments surrounded by a vanishingly low bulk actin concentration (Fig. 1A & B) (17). These droplets have properties of liquids, including an interfacial tension that determines their shape, growth through coalescence, and subunit diffusion (17). However, since the liquid is made of densely packed filaments, the filaments entropically align to form a nematic liquid crystal, where filaments have orientational order, giving rise to the tactoid shape (26, 27).

**Figure 1.**
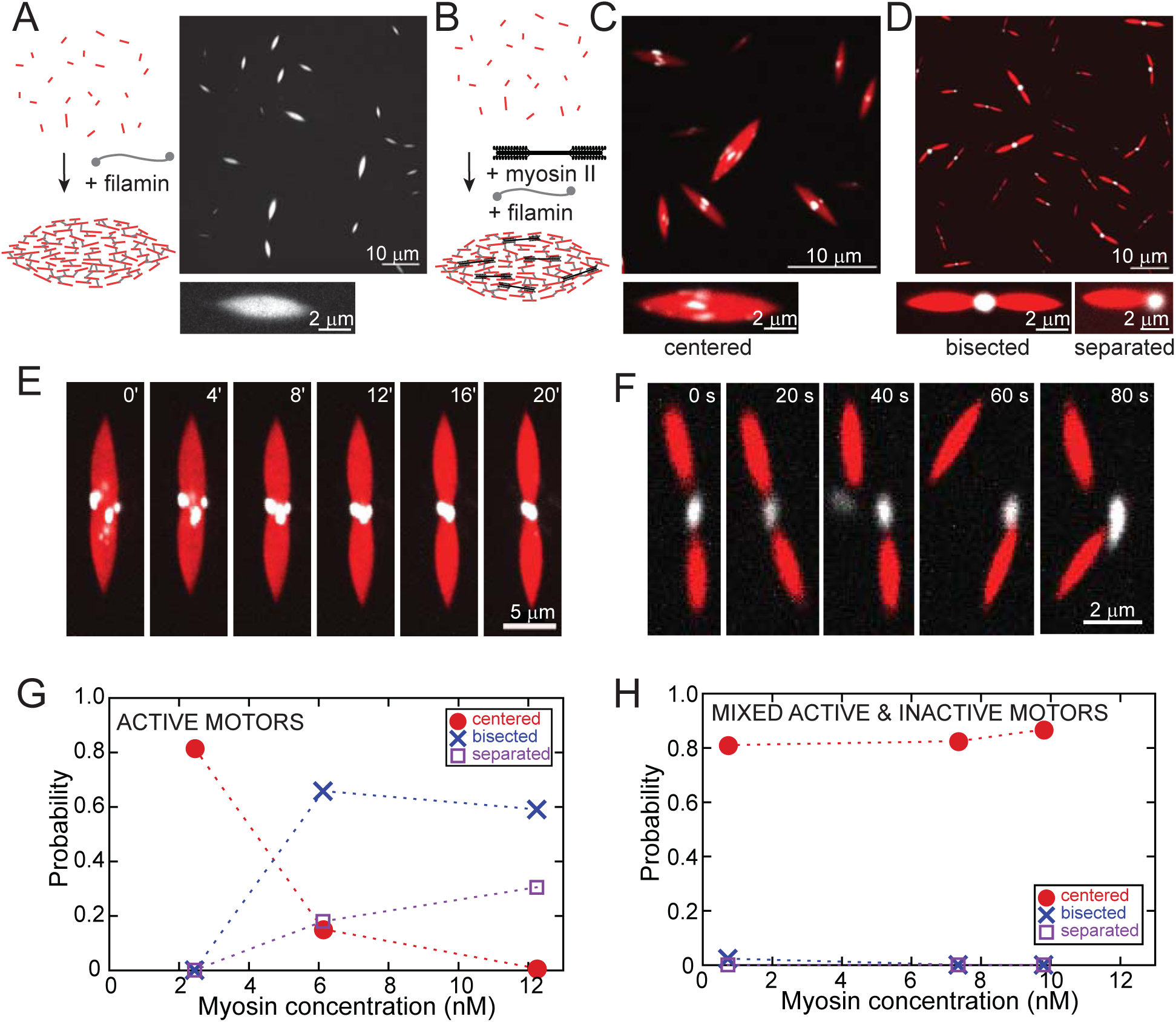
Molecular motors self-organize and deform biopolymer droplets. (A) Cartoon of experimental system used to make actin droplets. Fluorescence images of 10 mol% filamin cross-linked short actin filaments (2.6 μM actin, labeled with TMR) condensed into droplets. (B) Cartoon of experimental system used to make actin droplets containing myosin filaments (C) Upper: Image of myosin II (7.4 nM, labeled with Alexa 642, partially inactive, white) organized at the midplane of actin droplets (2.6 μM actin, red). Lower: close up of single droplet with motors “centered” (lower). (D) Upper: image of myosin II (12.3 nM, active, white) clustered at the center of “bisected” droplets (2.6 μM actin, red). Lower: close-up image of a bisected droplet, with the myosin cluster bridging droplets in a dimer and a “separated” droplet, where the myosin cluster localizes to the droplet pole. (E) Fluorescence microscopy images of myosin II puncta (0.7 nM, white) migrating to the actin droplet (2.6 μM, red) midplane as single rods, clustering, and bisecting the droplet (0 min defined as the beginning of data collection). (E) Image sequence of actin droplets (2.6 μM, red) severing from a motor cluster (12.3 nM myosin, active, white). (F) Distribution of actin-myosin droplet morphologies as a function of myosin concentration for active myosin motors, with fraction of “centered”, “bisected” and “separated” morphologies observed. (G) Distribution of actin-myosin droplet morphologies as a function of myosin concentration for samples that contain partially inactive myosin.

To investigate the effects of molecular motors, we create composite actin droplets containing skeletal muscle myosin II. Under these conditions, the myosin polymerizes into filaments, where each filament has hundreds of individual myosin heads and is ∼1 μm long, and ∼30 nm in diameter (28). As with pure actin droplets, adding the cross-linker filamin to a solution of pre-polymerized actin and myosin II induces the condensation of actin droplets embedded with myosin motors (Fig. 1B). These composite droplets have strikingly consistent features and shape, which are modulated by myosin concentration. For low motor concentration, we observe that myosin accumulates at the midplane of the droplet (Fig. 1C & Movie S1). The rod-like myosin puncta are near diffraction-limited in width and ∼0.3−1.6 μm in length, consistent with individual myosin filaments. The rod-like puncta align with the tactoid long-axis and stack across the droplet midplane. At higher myosin concentration, myosin II forms micrometer-sized clusters. These clusters localize to droplet poles, with the remaining droplets devoid of myosin (Fig. 1D & Movie S2). The myosin clusters either join two similarly-sized actin droplets together (“bisected”) or associate with isolated droplets (“separated”) (Fig. 1D & Movie S2).

We use timelapse fluorescence microscopy to explore the formation of bisected droplets. Although we miss the earliest droplet configuration, we find that isolated myosin puncta accumulate and cluster at the droplet midplane, over a few minutes (Fig. 1E and Movie S3, *t*=0 is defined as the beginning of the movie). As myosin clusters accumulate, they become larger and rounder and locally distort the actin droplet near the myosin cluster. This eventually leads to a single, micron-sized myosin cluster that distorts the droplet interface significantly enough to bisect the droplet into two smaller droplets (Fig. 1F). Bisected droplets are dynamic, where droplets occasionally spontaneously detach from the myosin cluster, yielding two individual, myosin-free droplets and an isolated myosin cluster over ∼1 min (Fig. 1F, Movie S4 & S5). Thus, actomyosin droplets exhibit spontaneous component centering and break up into two equal daughter droplets, reminiscent of active processes critical in biological cell division.

Composite droplet morphology is dependent on motor concentration. At the lowest concentrations (2.4 nM), motors exclusively center to the droplet midplane (Fig. 1G). As the motor concentration increases beyond 6 nM, motors cluster and primarily form bisected droplets, with more separated or single droplets at the highest motor concentration (Fig. 1G). To clarify the extent to which motor enzymatic activity is essential for these morphologies, we formed myosin filaments containing a proportion of inactive heads; that is, myosin that binds to actin without undergoing the power stroke (SI) (29). In these partially inactive myosin puncta, the myosin organizes at the droplet midplane over all myosin concentrations, without clustering in or dividing droplets (Fig 1H). This indicates that although enzymatic activity is necessary for droplet bisection and separation, it is not necessary for myosin centering.

## Myosin Filament and Droplet Structure Drives Midplane Localization

The earliest droplets that we observe already have myosin at the midplane. Nevertheless, we can investigate the centering process by examining droplets shortly after they diffuse into each other and coalesce into a larger droplet (17). Immediately after coalescence of composite droplets, myosin is distributed throughout the droplet (Fig. 2A, Movie S6). Over minutes, myosin puncta migrate to the midplane (Fig. 2A, Movie S6). The probability distribution of myosin localization as a function of distance, *d,* from the droplet midplane normalized to its length, *L*_D_, for 50 coalescence events indicates that this centered organization is general (Fig. 2B). Within two minutes of coalescence events, ∼90% of myosin puncta are dispersed along the central 40% of the droplet long axis (Fig. 2A, 2B). By 5 min, 65% of myosin puncta are at the midplane. Thus, myosin filaments initially located in distal droplet regions migrate to the droplet midplane.

**Figure 2.**
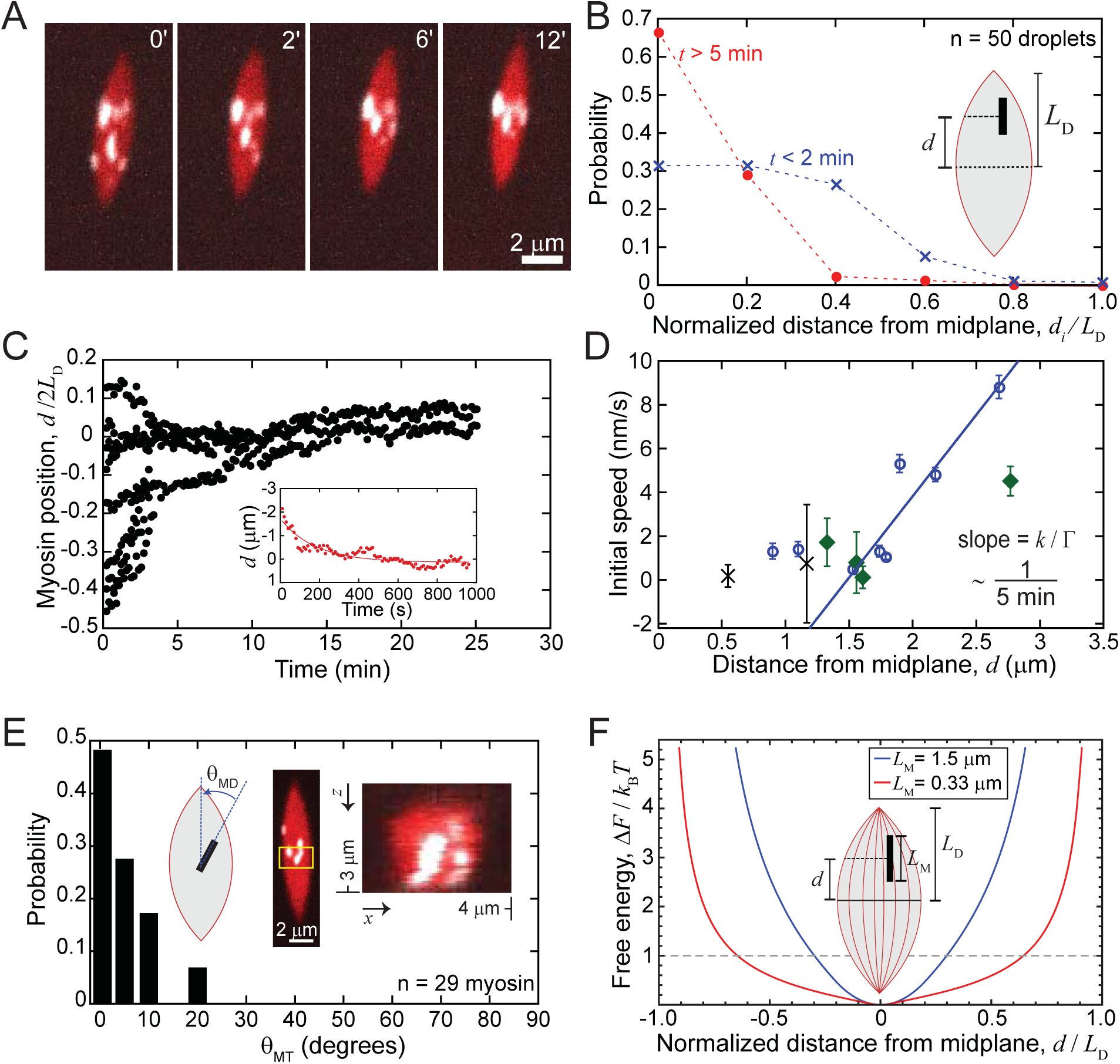
Motors migrate to regions of minimum nematic splay. (A) Fluorescence microscopy image sequence of myosin II motors (7.4 nM, partially inactive, white) in a cross-linked actin droplet (2.6 μM actin, red, where *t*=0 min is defined by when tactoid reaches rest length after coalescence, see supplementary movie). (B) Probability distribution for motors along the droplet long axis, binned at regular intervals as a function of distance from the droplet midplane, normalized to the droplet length *L*_D_ (n=50 droplets). (C) Myosin II puncta positions in the droplet in (A) over time. Distance is measured with respect droplet midplane. Lines represent exponential fits to the data. (D) Initial myosin II puncta speed as a function of initial distance from the droplet midplane. We observe that the centering speeds, v of myosin puncta more than a micron away from the droplet midplane scale linearly with distance from the midplane, d, as expected for a particle diffusing in a harmonic free energy landscape. The slope of the line of best fit to the v/d has a slope of 0.0032 *s*^-1^, which corresponds to a centering time of about 5 minutes. (E) Probability of alignment of myosin II puncta along droplet axis. Inset: fluorescence microscope images of myosin II puncta aligned at midplane. Averaging over the region in yellow in y, and projecting z indicates that myosin II puncta are distributed through the droplet bulk. (F) Model free energy plots of two myosin puncta of different lengths as a function of their position within a tactoid of length, *L_T_* = 3.2 *μm*. The free energy scale in relation to kT is determined from the diffusive kinetics of centering puncta obtained from Fig. 2D.

To gain insight into the mechanism of myosin self-organization, we track the dynamics individual myosin puncta. Tracking the normalized distance from the midplane, *d*/*L*_D_, as a function of time suggests that myosin puncta initially located far from the midplane have steep trajectories directed towards the midplane where *d*/*L*_D_=0 (Fig. 2C). By contrast, myosin initially located near the midplane undergo only small displacements (Fig. 2C). Taking the myosin speed to reflect the free energy landscape, these data suggest that myosin at the droplet midplane may be low energy compared to distal droplet locations. To quantify how myosin speed varies across the droplet, we extract the initial velocity from a linear fit to the early trajectory (Fig. 2C, inset, SI). Myosin puncta located near the midplane (*d* < 1.5 μm) have low initial speeds (∼1.4 nm/s), while their speeds increase to ∼ 6−10 nm/s when farther from the midplane (*d* > 2 μm) (Fig. 2D). These speeds are at least 100-fold slower than the unloaded gliding filament velocity of skeletal muscle myosin (30, 31). The low speeds indicate that actomyosin sliding is not primarily responsible for myosin centering, consistent with our finding that centering occurs with strongly bound, but not cycling, motors (Fig. 1H).

More than 90% of the centered myosin puncta align within 10° of parallel to the droplet major axis (Fig. 2E). This is consistent with the known preference of myosin II filaments to preferentially bind to actin filaments such that there is a high degree of alignment of their long axes (32). By sectioning the droplet midplane in *z*, we find that myosin are not only aligned but also embedded throughout the bulk of the droplet, indicative of strong actin-myosin interactions (Fig. 2E, inset). Actin filament orientation varies spatially across the liquid crystal droplet. Since we expect actin filaments to be splayed at the droplet poles and parallel to the droplet long axis near the midplane (Fig. 2F, cartoon, red lines indicate filament alignment) (33), we hypothesize that centering arises from the energetics of embedding the myosin rod in a structured actin droplet.

To gain further insight, we develop a continuum model where we model a single myosin II filament as an elongated rod of length, *L*_M_, that interacts with orientationally aligned actin filaments in its vicinity. The myosin rod is embedded in a droplet with a long axis, 2*L*_D_, where a bipolar director configuration represents the local alignment of actin filaments. In this configuration, the curved director lines are parallel at the midplane and splayed near the droplet poles (Fig. 2F, cartoon, SI, Fig. S1). Myosin II filaments are ∼30 nm wide, thin enough not to distort the surrounding actin nematic, in which we estimate a ∼50 nm spacing between actin filaments (17). We impose a strong orientational preference to represent the mutual alignment of myosin and actin filaments (32). Based on this model geometry, we calculate the energetic cost of misalignment of the myosin rod, Δ*F*(*d*), expressed in terms of rod displacement, *d*, from the droplet midplane (SI). The resulting free energy landscape of the rod has a minimum at the droplet midplane (Fig. 2F). As *d* increases, the rod interacts with more splayed directors resulting in larger misorientation angles and higher energy costs. For a longer rod (*L*_M_=1.5 μm, Fig. 2F, blue), the cost of misalignment is higher than for a short rod (*L*_M_=0.3 μm, Fig. 2F, red).

To relate the model energy scale to thermal energy, we suppose the motors to be diffusive in the free energy landscape, Δ*F*(*d*). We consider that a single motor filament experiences centering force, *f*(*d*), and use a harmonic approximation such that *f*(*d*) *= kd,* where *k* is the spring constant. The associated free energy Δ*F = kd*^*2*^*/*2 may be written as 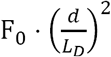, where *F*_0_ is approximately the work required to move a motor across the droplet. The centering speed, *v*, is determined by the equality of *f*(*d*) and the drag force, Γ*v*. The slope *v/d* ∼ 1/(5 min) (Fig. 2D) is the inverse centering timescale which equals *k*/Γ. From the previously estimated 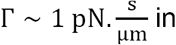, actin tactoids (17), we extract *k* from the slope. Using the corresponding energy scale, 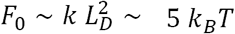 (SI), we rescale the free energy values with respect to thermal energy (Fig. 2F). The model predicts that the motor filaments drift towards the droplet midplane and center up to distances at which the centering energy becomes comparable to the thermal energy. We expect low centering speeds within a few microns of the midplane (Fig. 2F), consistent with experimental measurements of low velocities seemingly independent of *d* in that region (Fig. 2D). Thus, rod-like myosin filaments center by optimizing their location within the structured actin droplet.

## Myosin clusters deform and divide droplets through disrupting nematic aligment

We next sought to understand how myosin activity at the droplet midplane results in droplet deformation. In the absence of myosin activity, actin droplets do not spontaneously divide (17), suggesting that myosin activity generates forces that counteract droplet interfacial tension and actin alignment to drive shape changes. A feature of active myosin is that it forms clusters, so we inspected myosin clusters of different size and shape in droplets. Rod-like myosin filaments embedded within the actin droplet cause no local shape distortion or variation in the actin intensity (Fig. 3A(i)). With active myosin, multiple myosin filaments aggregate into micron-sized clusters. During clustering, puncta increase in area and become rounder (Fig. 3A(ii-iv). Simultaneously, the droplet shape distorts at the myosin cluster location. Near large clusters, the amount of actin is depleted, as a precursor to droplet separation (Fig. 3A(iv)).

**Figure 3.**
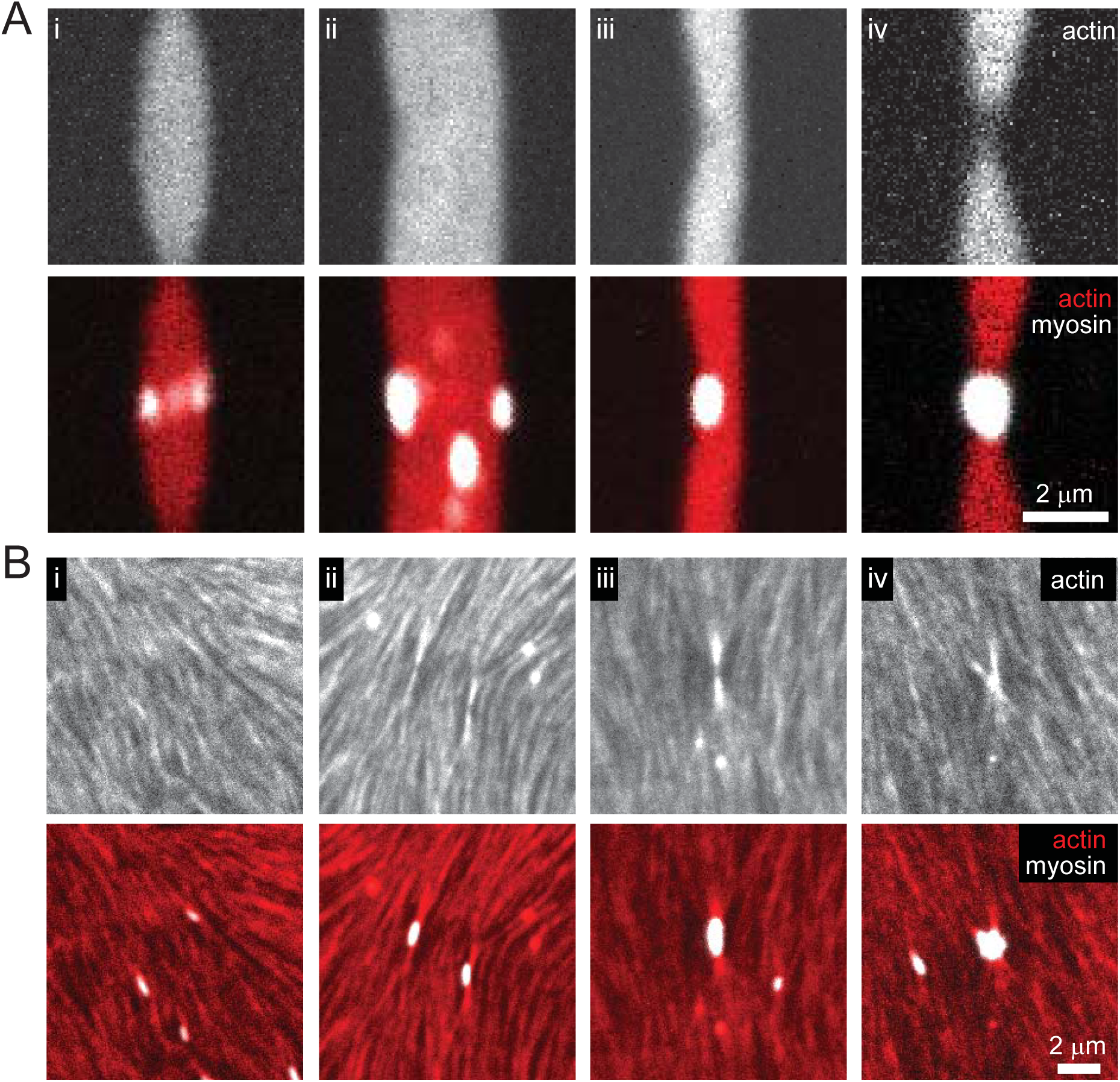
Molecular motors distort actin alignment and induce droplet deformations. (A) Fluorescence images of actin (2.6 μM, upper) and actin and myosin (lower, actin is red and myosin is white) in composite droplets. Myosin puncta do not visibly distort actin droplets (column 1), while small myosin clusters distort the droplet interface (column 2 and 3), and large clusters divide droplets (column 4). (B) Fluorescence images of a thin actin nematic (2.6 μM actin, 1.2 mol% capping protein, crowded with 0.4 wt. % methylcellulose), showing actin (upper row, lower red) and myosin (1.1 nM myosin, lower, white).

We propose that droplet distortions arise from changes in actin filament organization around the myosin during enzyme activity-mediated clustering. Colloidal inclusions suspended in a nematic liquid crystal can distort the surrounding nematic (20). The nature of the distortion induced by a colloid depends on its size and anchoring, the preferential alignment of liquid crystal constituents at the colloid surface. To investigate this, we examine the impact of myosin cluster shape and size on actin filament alignment in thin, bulk nematic actin liquid crystals, where actin alignment is visible (SI) (23). In bulk nematics, rod-like myosin puncta align parallel to actin filaments (Fig. 3B(i)). By contrast, myosin clusters impose local distortions in the nematic, such that actin filaments are canted around the cluster (Fig. 3B(ii-iv)), consistent with previous data showing that myosin clusters impose a radial arrangement of actin filaments (34).

Motivated by these experimental observations, we extended our continuum model to capture dividing droplets by representing the myosin cluster as a spherical colloid that imposes anchoring normal to its surface in a nematic droplet. In a bulk nematic, a colloid with normal anchoring creates defects in its vicinity (20, 21). In contrast to typical colloidal inclusions, myosin clusters bind to the surrounding actin, which we represent by an adhesion energy per unit area on the colloid surface, *w*. The bipolar droplet shape results from an optimization of the bulk nematic elastic and interfacial energy, with strong anchoring at the droplet interface (33). Since the myosin clusters impose normal anchoring, the radial convergence of the actin filaments near the droplet pole presents an optimal anchoring configuration. Like any liquid droplet, a single nematic droplet has lower interfacial energy than two such smaller droplets with the same total volume. However, we find that for droplets in contact with an adhesive colloid, the energetic gain from increased wetting of the colloid’s surface counteracts the elastic and interfacial energetic cost of deforming the droplet, leading to the minimum configuration of a bisected droplet (SI). As the colloid size or adhesion strength increases compared to the droplet interfacial tension*, w/γ*, the resulting adhesive energy gain overwhelms the energetic cost of increasing the interface through shape distortions. In this case, two new droplet poles are created and the motor cluster positions at the center of two, smaller droplets (Fig. 4C, blue). In contrast, when the adhesion is small compared to the interfacial tension, the energetic minimum occurs when the colloid is at a droplet pole (Fig. 4C, red), consistent with observations of colloids in nematic molecular liquid crystal droplets (35).

**Figure 4.**
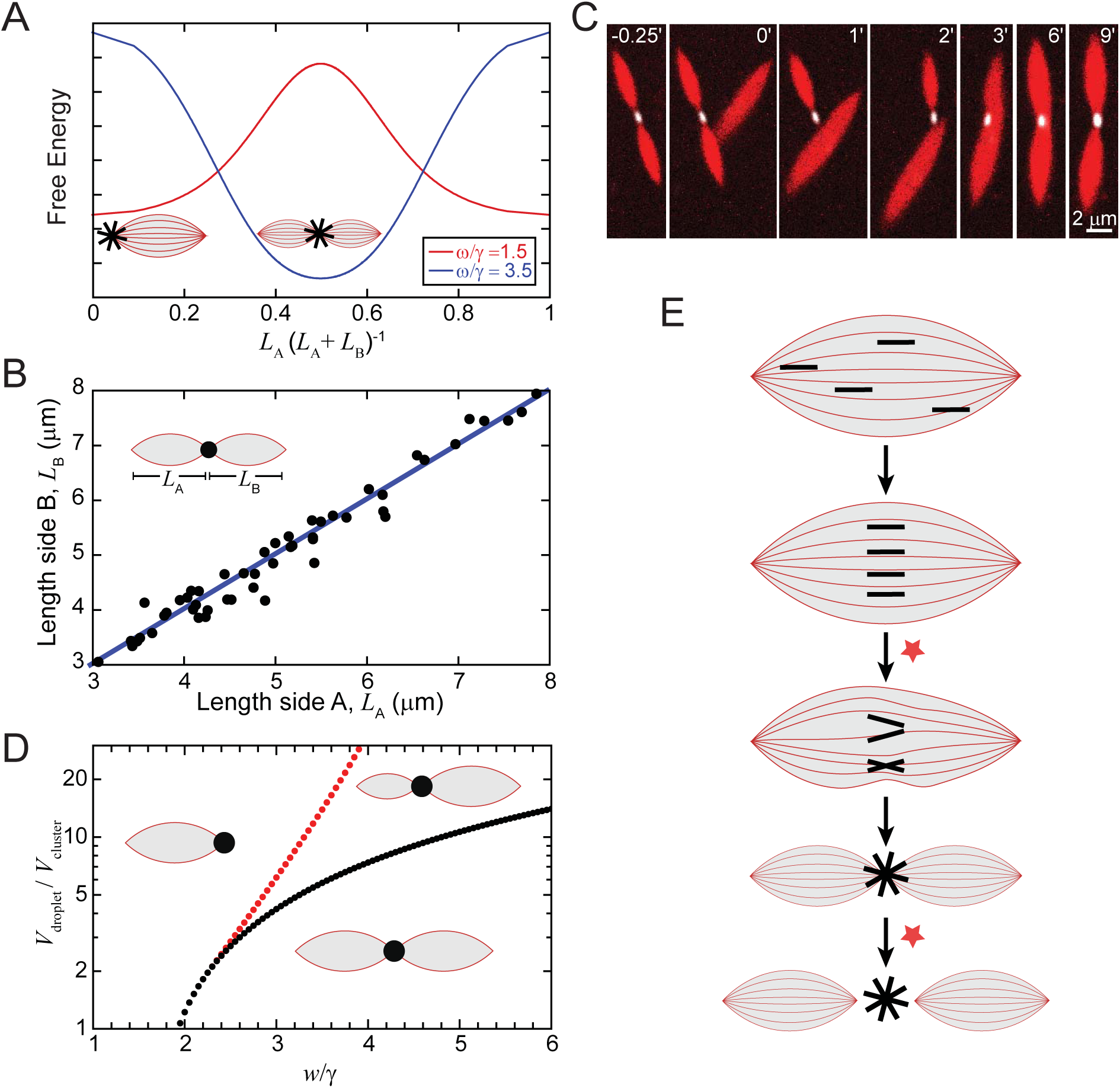
Molecular motors bisect biopolymer droplets. (A) Model free energy plots show that droplet division can be favored in the presence of an adhesive colloidal inclusion. The total free energy of two divided nematic bipolar droplets with lengths, *L*_*A*_ and *L*_*B*_, of fixed total volume, that adhere to the motor cluster modeled as a colloid with normal anchoring and adhesion energy *w* per unit area, is shown as a function of a droplet division parameter: *L*_*A*_ · (*L*_*A*_ + *L*_*B*_)^−1^. The mechanical energy of each droplet includes contributions from interfacial energy given by surface tension, *γ*, and nematic elastic energy. At higher values of *w* compared to a relevant droplet mechanical energy parameter, the equal division of droplets becomes favorable (blue curve) as opposed to a single whole droplet which is the preferred state for low *w* (red curve). The free energy calculated here is for nearly spherical bipolar tactoids. The complementary case of very elongated bipolar tactoids is shown in the SI and has the same qualitative behavior with equal division favored at higher adhesion. Both free energy curves have been normalized to display their minima. (B) Comparison of the length of one side of a divided droplet to the other. Blue line indicates a slope of one. (B) Fluorescence images of a droplet coalescing with a bisected droplet (2.6 μM actin, 12.3 nM active myosin). (D) Phase space predicted by continuum model of droplet division resulting from competition of droplet mechanical energy and adhesion with colloid. At higher values of adhesion, w, and lower droplet volume in relation to the colloidal motor cluster, division into two equal droplets is energetically favored. The black dots indicate where the equally divided state changes to a free energy minimum from a maximum (see Fig. 4A) as adhesion is increased. As adhesion decreases (or correspondingly droplet volume increases), unequal division is favored. At the red line, the smaller unequal droplet becomes much smaller than the other droplet in the favored configuration, indicating that a single whole droplet is favored at lower adhesion. (E) Cartoon summarizing motors in biopolymer droplets. Red stars indicate transitions that activity is necessary to complete.

## Droplets Prefer to Divide Equally

Strikingly, myosin clusters typically bisect droplets, with the resultant two droplets always apparently equal in size (Fig. 2B). Indeed, comparing the lengths on either side of the dimer indicates that the motor cluster precisely divides the droplets into two equal droplets (Fig. 4B). The coalescence of a divided droplet dimer with an isolated droplet provides further evidence for the strong preference for droplet bisection (Fig. 4C, Movie S7). In this case, an isolated droplet coalesces with one droplet of the dimer, temporarily leading to the myosin cluster bridging two droplets of disparate sizes (Fig. 4C, 1-2 min). However, the actin redistributes over a few minutes until the droplets on either side of the myosin cluster are again equal. This indicates that the dimer with equal-sized droplets is an energetically favored rather than a trapped configuration, that might have resulted from the individual myosin puncta migrating to the droplet midplane prior to forming a cluster. Thus, the net adhesive energy gain is large enough to favor redistribution of droplet material into two equal droplets. We can theoretically explore how systematically changing the adhesion influences the preferred configuration of the actin droplet and myosin cluster. For low adhesion, the preferred configuration is a single large droplet with a polar myosin cluster. At intermediate adhesion, two droplets of unequal volume can form as a result of the competition between droplet interfacial energy and adhesion with colloid surface, while at higher adhesion the two droplets are of equal volume (Fig. 4D).

## Discussion

Here, we have introduced structured fluid droplets that exploit geometry sensing to generate spatial organization and drive shape change. While there are many potential manifestations of geometry sensing, we focus on droplets that exhibit essential aspects of cell division: center-finding and division into two identical daughters. The droplet shape evokes the mitotic spindle and the center-finding is reminiscent of chromosome alignment at the spindle midplane. That our droplets are comprised of biomolecules quite distinct from those in the spindle, highlights potentially shared physical mechanisms. In fact, spindle shape and dynamics is captured by the physics of active structured fluids (25). While structured fluids, such as molecular liquid crystals, have long been used to organize colloids at defects and interfaces (20, 21, 35-38), here we exploit colloid affinity to the liquid constituents to change the energetically favored locations within the liquid crystal bulk. This interplay of adhesion, interfacial tension, and elasticity potentially has implications in regulation of spatial organization in sub-cellular liquid droplets beyond the spindle, in primitive and synthetic cells, and other macromolecular droplets such as coacervates (25, 39-41). Intriguingly, we find these processes, including center finding, do not rely on enzymatic activity in the droplet or colloid, but rather that the colloid strongly interacts with the droplet. In the future, this system can be extended to elucidate how colloid adhesion and shape, together with fluid structure, direct spatiotemporal organization in structured fluids, which may inform the design of novel structured, soft materials.

A key feature of biological cells that remains elusive to reconstruct in synthetic systems is cell division (41, 42). Driving droplet shape changes has an energetic penalty associated with increasing the amount of droplet interface due to an interfacial tension. Typically, activity is invoked in theoretical arguments to provide the energy associated with deforming the interface (16, 43). By contrast, here we demonstrate a system where droplet division occurs passively, as a result of the competition between colloid adhesion with the droplet and the droplet energetics, while activity serves only to regulate changes in colloid shape and binding interactions. While the fluid structure in molecular liquid crystals is used to spatially organize colloids in droplets (21, 35, 38), to our knowledge, using colloidal wetting is a new approach to drive droplet deformations. In particular, we present a robust approach to drive droplet division into two equal daughters. In lyotropic systems such as these actin-based droplets, the entropic effects are much more prominent as compared to molecular liquid crystals due to the longer lengths of the nematogens (actin filaments) and colloids (myosin filaments). This different length scale may be critical to using colloids to drive shape changes. An exciting area of future research will be to utilize large, macromolecular liquids and chemical activity based on these mechanisms to sense geometry, robustly organize, and induce deformations to create novel, adaptive soft materials.

## Acknowledgements

We acknowledge T. Thoresen and S. Stam for purified filamin and C. Suarez for advising the purification of capping protein. We thank J. Vieregg, D. Coursault, R. Zhang, and P. van der Schoot for inspiring and insightful discussions. This research was primarily supported by the University of Chicago Materials Research Science and Engineering Center (NSF DMR grant 1420709). Additional support from the Sloan fellowship to S.V., ARO MURI grant W911NF1410403, NSF 1344203 and NIH GM085087 to M.L.G.

